# Functional independence of endogenous μ- and δ-opioid receptors co-expressed in cholinergic interneurons

**DOI:** 10.1101/2021.04.23.441175

**Authors:** Seksiri Arttamangkul, Emily J. Platt, James Carroll, David L. Farrens

## Abstract

Class A G protein-coupled receptors (GPCRs) normally function as monomers, although evidence from heterologous expression systems suggests they may form homodimers and/or heterodimers. Detection of GPCR dimers in native tissues has however been challenging due to the lack of suitable tools. μ- and δ-Opioid receptors (MORs and DORs) co-expressed in transfected cells has been reported to form heterodimers. The co-localization of MORs and DORs in neurons has been studied in knock-in mice expressing genetically engineered receptors fused to fluorescent proteins. Here we report that single cholinergic neurons in the mouse striatum endogenously express both MORs and DORs. The receptors were fluorescently labeled in live brain slices with a ligand-directed labeling reagent, NAI-A594. The selective activation of MORs and DORs, with DAMGO (μ-agonist) and deltorphin (δ-agonist) inhibited spontaneous firing in all cells examined. In the continued presence of agonist, the firing rate returned to baseline with the application of deltorphin but was persistently inhibited with the application of DAMGO. In addition, agonist-induced internalization of DORs but not MORs was detected. When MORs and DORs were activated simultaneously with [Met^5^]enkephalin, desensitization of MORs was facilitated but internalization was not increased. Together, these results indicate that while MORs and DORs are expressed in single striatal cholinergic interneurons, the two receptors function independently.

## Introduction

Opioid receptors are members of GPCRs targeted by endogenous peptides and exogenous opiate drugs. Three opioid receptor-subtype, μ (MOR), δ (DOR) and κ (KOR) are known to be distributed in many brain areas (Le Merrer et al., 2009). The co-expression of opioid receptor-subtypes in individual neurons has steadily gained attention because possible cross-interactions between the receptors may serve as new pharmacological targets for analgesia while avoiding the side effects resulting from activation of a single opioid receptor subtype (Fujita et al., 2015). Although some biochemical and fluorescent imaging studies support the existence of MOR-DOR heterodimers (Cahill and Ong 2018), the findings required the use of heterologous expression and genetically modified receptors to enable experimental detection. Thus, the key evidence that both MORs and DORs are endogenously co-localized in single neurons has been questioned. The major challenge is that opioid receptors are generally expressed in low densities in neuronal tissues thus detection of subcellular co-localization is often ambiguous. Functional readout using electrophysiological measurement is a powerful tool to detect MOR-DOR co-expression in single neurons but there are few such studies (Egan and North, 1981; Chieng et al., 2006).

The striatum is a subcortical structure in the forebrain that plays an important role in motivation, sensorimotor function, goal-directed learning and drug addiction (Kreitzer 2009; Castro and Bruchas 2019). The largest portion of neurons are medium spiny GABAergic projecting neurons that are also known to produce opioid peptides including enkephalins (μ-and δ-agonists) and dynorphins (κ-agonists). The striatum consists of a heterogeneous structure where all opioid receptor-subtypes have been located by various techniques (Le Merrer et al., 2009). MOR binding sites are primarily detected in the striosomal patches whereas DOR and KOR binding sites are found in both matrix and patches (Herkenham and Pert, 1981; Tempel and Zukin, 1987; Kitchen et al., 1997; Banghart et al., 2015). Cholinergic interneurons (ChIs) comprise only a small population (1-2% of all neurons) and are scattered throughout the striatum. These neurons function as an important regulator of the synaptic activity within the striatum including the local release of dopamine (Cai et al., 2021; Yorgason et al., 2017; Threlfell et al., 2012; Cachope et al., 2012). ChIs are spontaneously active neurons that constantly fire action potentials and release acetylcholine in brain slices and *in vivo* (Wilson et al., 1990; Bennett and Wilson 1999; Zhou et al., 2002). The firing rate of these neurons is inhibited with the activation of opioid receptors. Jiang and North (1992) recorded hyperpolarization of the secondary neurons (presumably ChIs) and reported that these neurons were sensitive to δ- but not μ-agonists. Other studies however report the presence of functional MORs in these neurons (Jabourian et al., 2005; Britt and McGehee, 2008; Ponterio et al. 2013; Mamaligas et al., 2016). It is possible that various ChI subpopulations contain segregated MORs and DORs (Jabourian et al., 2005; Britt and McGehee, 2008; Laurent et al., 2012). To date, there is no evidence demonstrating that MORs and DORs might be co-localizing in ChIs.

In the present study, we used image analysis and electrophysiological techniques to investigate the distribution of MORs and DORs on ChI neurons in the striatum. Live-cell imaging using NAI-A594, revealed fluorescent labeling of both MORs and DORs on the plasma membrane of single ChIs in brain slice preparations. The selective antagonists at DOR (SDM25N) or MOR (CTAP) were used to differentiate the two receptors and assess the degree of MOR and DOR expression. The inhibition of spontaneous firing using the selective μ-agonist (DAMGO) and δ-agonist (deltorphin II) confirmed that both MORs and DORs were present and functional on single neurons. [Met^5^]enkephalin (ME) and morphine also inhibited the firing activity of ChIs. As expected, ME interacted with both receptors while morphine selectively activated MORs. Interestingly, while the extracellular recordings showed activation at each receptor fully inhibited firing activity of the neuron, the agonists only induced substantial desensitization of DORs, not MORs. Similarly, only the DORs showed agonist-induced receptor internalization. Co-activation of the receptors by exogenous application of ME increased MOR desensitization. Collectively, the results in this study indicate that co-localized MOR-DOR function autonomously and there is little evidence for stable heterodimer formation. Nonetheless, downstream functional interactions may occur when both receptors are simultaneously activated.

## Results

### Ligand-directed labeling of opioid receptors

Naltrexamine-acylimidazole-alexa594 (NAI-A594; Figure 1A-a) is a labeling reagent that has been shown to specifically bind and then covalently tag the fluorophore Alexa-594 to MORs in brain slices (Arttamangkul et al., 2019). The labeling of this molecule is based on traceless labeling affinity approach (Hayashi and Hamachi, 2012; Shiraiwa et al., 2020), in which the naltrexamine moiety acts as a ligand that guides specific binding to opioid receptors. Once bound, the fluorophore is transferred to the receptor by the reaction of the acylimidazole group with an amino acid nucleophile, and at the same time, the guide ligand (naltrexamine) is cleaved and released from the binding pocket (Figure 1A-b).

**Figure 1.**
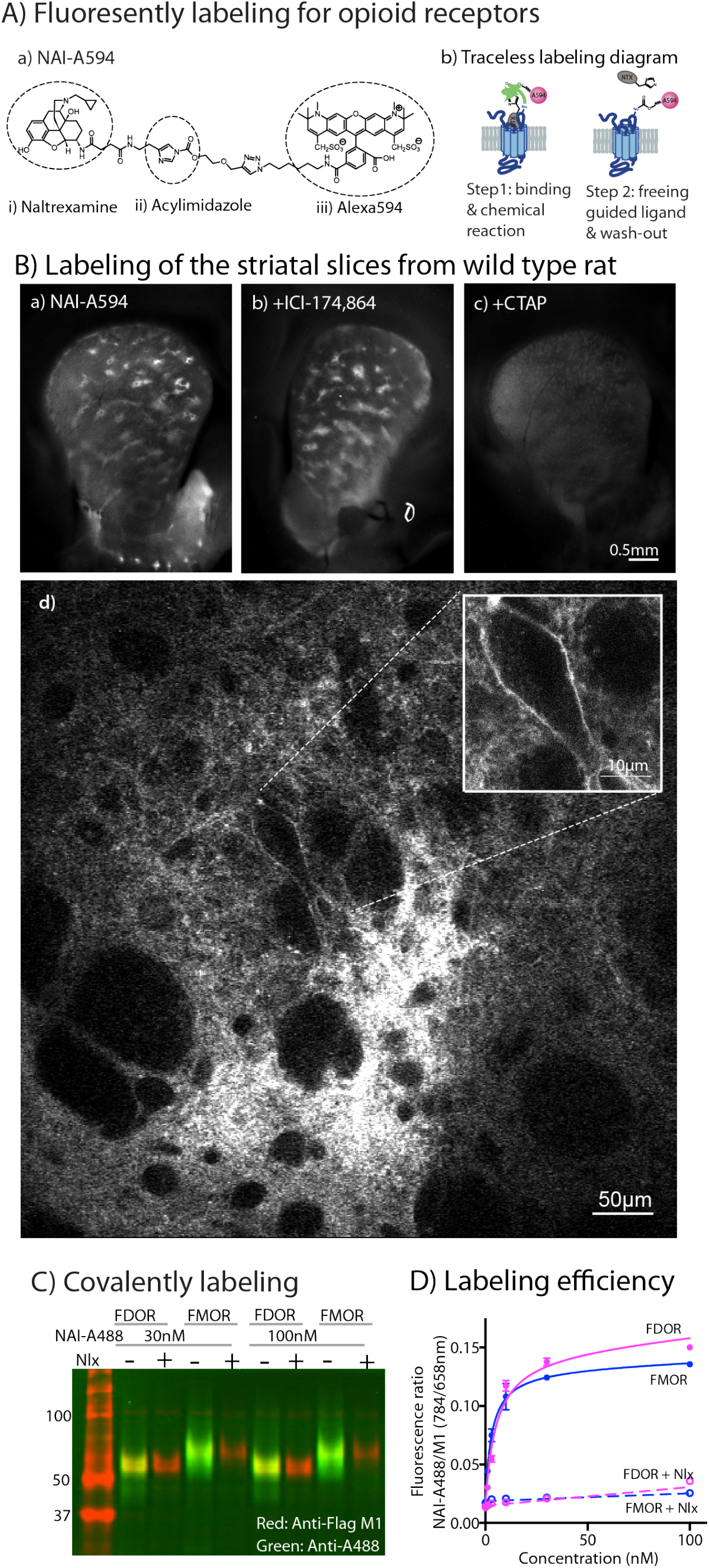
**(A)** Covalent labeling of opioid receptors with NAI compounds. (a) Chemical structure of naltrexamine-acylimidazole-alexa594 (NAI-A594). (b) Diagram of traceless labeling shown in 2 steps. Step 1, NAI compound binds and reacts with the receptor. Step 2, washing of naltrexamine moiety released from the reaction. **(B)** Live images of rat striatum incubated for 1 hour in (a) 100 nM NAI-A594, (b) 100 nM NAI-A594 plus 1 μM ICI-174,864, (c) 100 nM NAI-A594 plus 1 μM CTAP, and (d) zoom area of patch-like structure showing a large neuron believed to be the cholinergic interneuron at the boundary of the patch. **(C)** Near infrared western blots of HEK293 cells expressing FDOR and FMOR labeled with 30 and 100 nM NAI-A488 in the absence and presence of 10 μM naloxone, shown in a merge channel. The FLAG-epitope on opioid receptors was detected with anti-FLAG M1 (red), and the NAI-A488 modification of the receptors was identified with rabbit anti-Alexa Fluor 488 antibody (green). **(D)** Concentration dependent curves of FDOR and FMOR labeled with NAI-A488. The fluorescence intensity ratios of NAI-A488 (784 nm of secondary antibody to IR dye CW800) over anti-Flag M1 (658nm of secondary antibody conjugated to Alexa680) were plotted against NAI-A488 concentrations. The graph shows labeling curves performed in triplicate from one of three experiments. Data are shown as mean±SEM.

The labeling of striatal slices from wild type rats with NAI-A594 strikingly revealed striosomal patches, regions that are known to contain abundant MORs (Figure 1B-a and Arttamangkul et al., 2019). The MOR binding sites in the patches persisted when DORs were blocked with a selective antagonist ICI-174,864 (Figure B-b). In contrast, fluorescent labeling of the whole striatum was greatly diminished when the MOR selective antagonist, CTAP was included in NAI-A594 solution (Figure 1B-c). These results suggest that most of the labeling in the striatum were at MORs. Unexpectedly, some large neurons with a few dendrites near the boundary of the striosomal patches were also labeled (Figure 1B-d). The cell morphology and location of these neurons suggested that they were most likely ChI neurons (Brimblecombe and Cragg, 2016). The labeled receptors on these large neurons however could be either MORs or DORs because NAI-A594 was shown previously to stain Flag-tagged MORs (FMOR) or DORs (FDOR) in HEK293 cells (Arttamangkul et al., 2019).

To verify the comparable labeling of MORs and DORs with NAI-A594, a series of biochemical experiments were done using FMOR and FDOR cells. In these experiments, we used naltrexamine-acylimidazole-alexa488 (NAI-A488), which differed only the fluorophore from NAI-A594 so that we could detect the labeled receptors with an anti-Alexa488 antibody in Western blot analysis and compare the results to total Flag-tagged receptors using an anti-Flag antibody. The results showed that NAI-A488 at 30 and 100 nM reacted similarly with both receptors (Figure 1C, shown as yellow bands in a merge image). The labeled receptors and Flag-tagged receptors were clearly observed in the single channel images (Supplemental Figure 1A, green and red bands, respectively). The labeling of FDOR and FMOR by NAI-A488 was specific, as it could be completely blocked by addition of the universal opioid antagonist naloxone (Figure 1C, only red bands were shown). The fact that Alexa488-labeled receptors are detected even after denaturing gel electrophoresis confirms that the fluorophore is covalently bound to the receptors. The ability of NAI-A488 to label FDOR and FMOR was then analyzed by comparing concentration labeling curves of each receptor using an on-cell western assay (Figure 1D and Supplemental Figure 1B). The labeling curves for NAI-A488 reacted FDORs and FMORs were similar and saturated at about 100 nM. The labeling coefficients were also comparable, with apparent Kd values of 7.1 ± 2.8 nM for FDOR and 3.3 ± 0.7 nM for FMOR (unpaired t-test, p = 0.2543, n=3 experiments performed in triplicate). We conclude that the fluorescent NAI compounds can effectively and covalently tag dyes to MORs and DORs.

### Labeling of MORs and DORs on cholinergic interneurons with NAI-A594

To determine which opioid receptors are present on the striatal ChIs, we used brain slices from transgenic ChAT(BAC)-eGFP mice to identify ChIs with green fluorescent protein (GFP). Patch-like structures were similarly observed in the striatum from ChAT-GFP mice following incubation with NAI-A594 as shown earlier in rat (Figure 2A). The ChAT-GFP neurons were found scattered throughout the striatum (Figure 2B). The plasma membrane of GFP-positive perikarya, dendrites and axon terminals were also Alexa 594 positive (Figure 2C&E for NAI-A594 staining and D&F for the merged images of NAI-A594 labeling and GFP). There are neurons lacking GFP that also reacted with NAI-A594 (Figure 2C&D).

**Figure 2.**
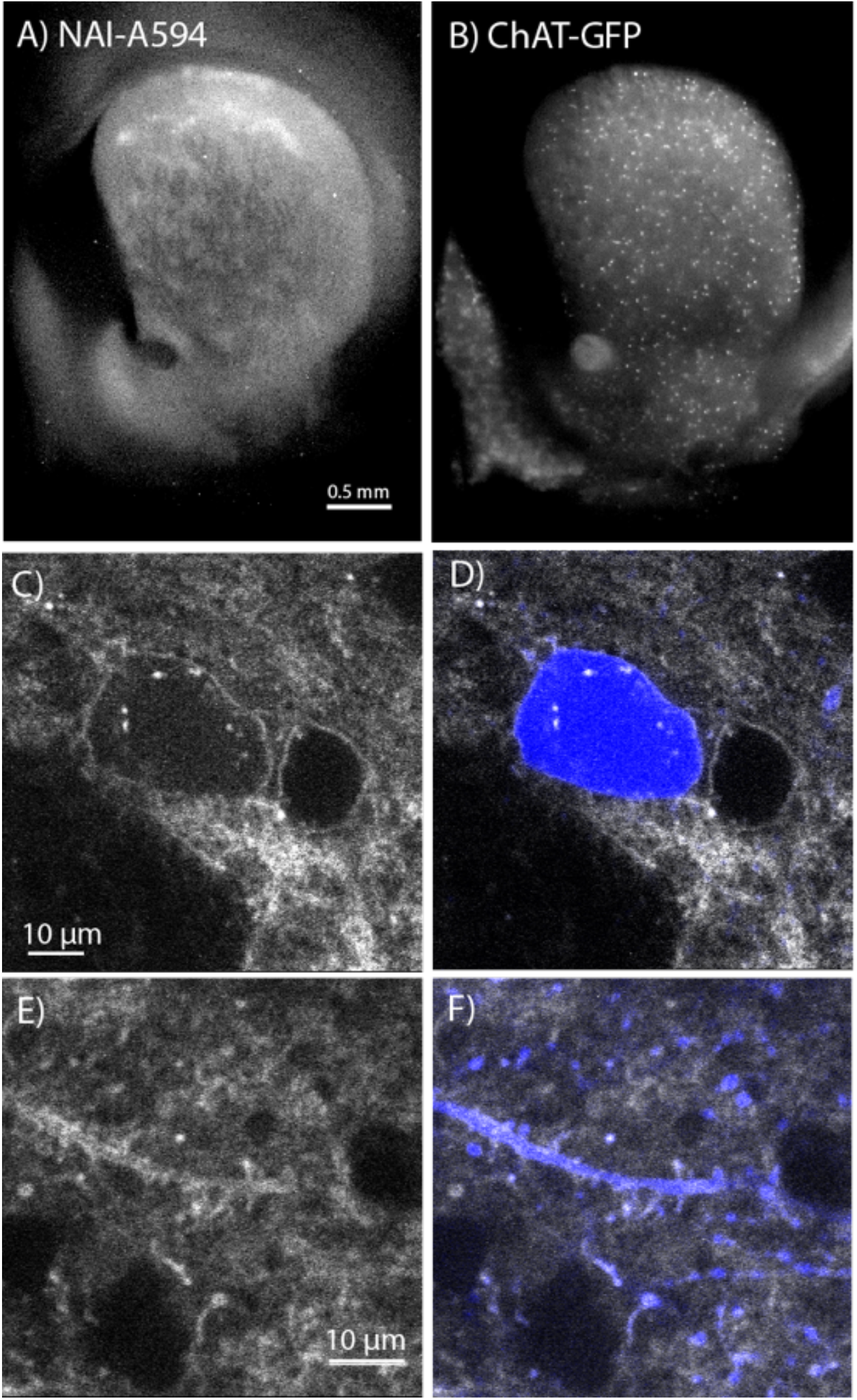
*In situ* NAI-A594 labeling of ChI neurons in the striatum of ChAT-GFP mice. **(A)** An area where a fluorescent patch is clearly observed in the striatum of ChAT-GFP after the staining of NAI-A594. **(B)** Distribution of GFP-positive neurons in the same section of brain slice in (A). **(C-F)** The labeling of NAI-A594 is clearly observed along plasma membrane staining of cell body, dendrites and terminals. The gray outline of two cells in (C) and the merged channel shows GFP (blue) in (D) indicate that the other neuron, presumably MSN is also stained with NAI-A594. Neurites containing GFP are visible after staining (E). Fluorescence of neurites are observed throughout the area. The merged image (F) shows NAI-A594 labeled both positive and negative GFP neurites.

Live-cell imaging experiments revealed all GFP-positive neurons reacted with NAI-A594 (n=246 from 7 male and 6 female mice). Figure 3A shows example images of GFP-positive cells labeled with NAI-A594 alone, NAI-A594 in the presence of μ- selective antagonist CTAP (1 μM) such that DORs alone were identified. In experiments with NAI-A594 in the presence of δ-selective antagonist SDM25N (0.5 μM), MORs were specifically detected. Finally with the incubation of NAI-A594 with both antagonists was used to block both MORs and DORs. The results show a partial decrease of the fluorescence on the plasma membrane of CTAP-treated neurons as compared to experiments with NAI-A594 alone (Figure 3A-b and -a). In the presence of SDM25N, the fluorescence staining was further reduced as compared to CTAP-treated cells (Figure 3A-c and -b). The combination of SDM25N and CTAP completely blocked the labeling of ChAT-GFP neurons (Figure 3A-d).

**Figure 3.**
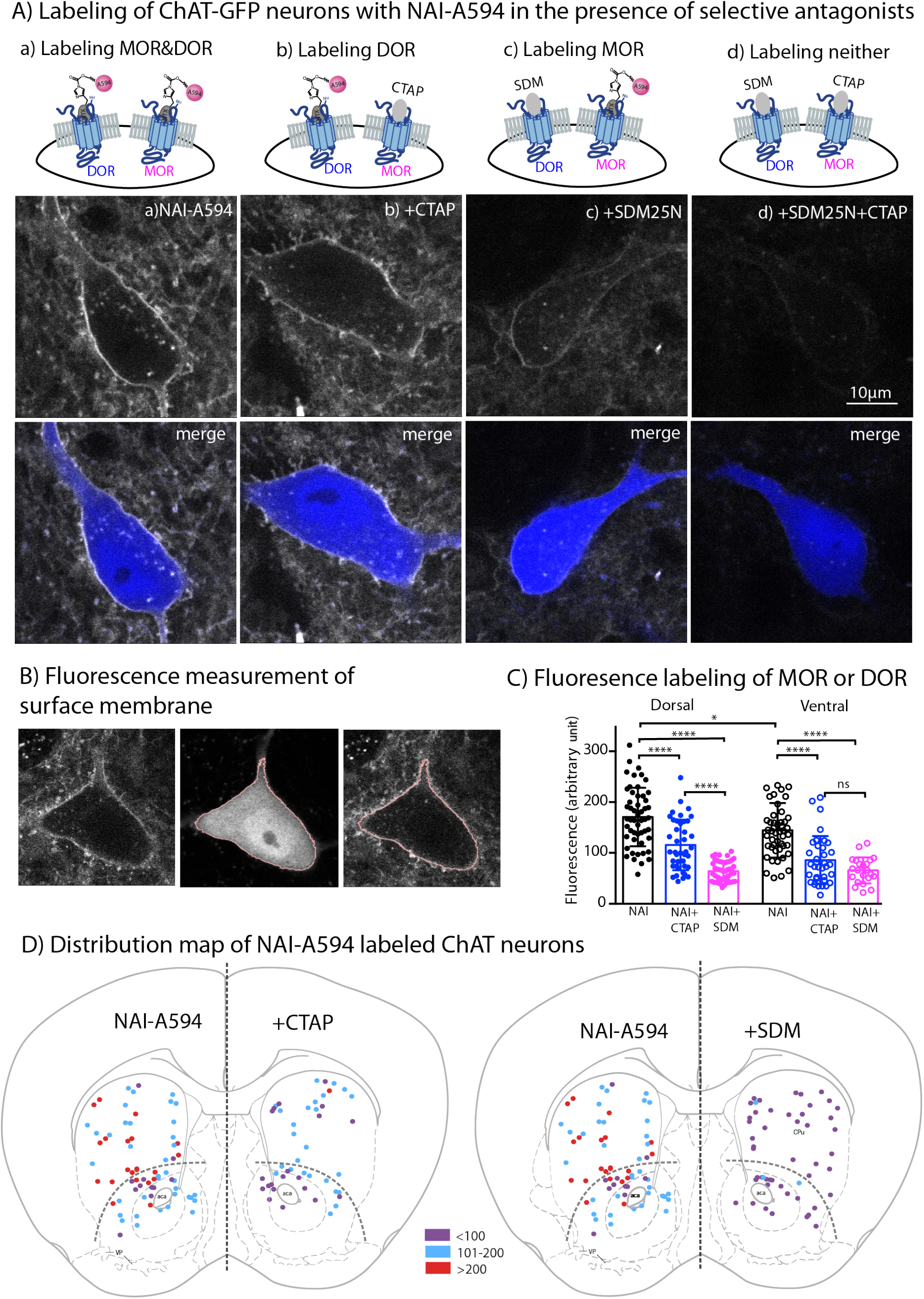
Fluorescent intensities of NAI-A594 labeling of endogenous MOR or DOR receptors. **(A)** Examples of single ChAT-GFP positive cell showing staining of NAI-A594 under different conditions: (a) NAI-A594, (b), NAI-A594+CTAP, (c) NAI-A594+ SDM25N, and (d) NAI-A594 +CTAP and SDM25N. The upper panel are images from Alexa594 channel, and the bottom panel are the merged images of GFP (blue) and Alexa594 (gray). **(B)** Diagram images illustrate the method of measurement of surface membrane fluorescent intensity. Left is the raw signal of a neuron labeled with NAI-A594. Middle shows outline boundary derived from GFP signal and thresholding using ImageJ (details in Materials and Methods). Right, the fluorescent intensities of Alexa594 channel are measured along the outline drawn in the GFP channel and supper-imposed onto the raw image. The lines drawn here are enlarged for illustration. The true thickness of line is ~0.08 μm or 1 pixel. **(C)** Summarized results of fluorescent intensities in each labeling condition. Dorsal and ventral areas are defined according to the mouse brain atlas (Franklin and Paxinos, 2007). All data are shown in mean±SD, p<0.0001 (****) and p<0.05 (*). **(D)** Distribution maps of the observed ChIs, in which fluorescent intensities were measured in different conditions. Data are presented as combined results from both male and female. Color codes represent the range of fluorescent intensities (in Arbitrary Units, AU) shown in (C).

To estimate the relative amount of MORs and DORs on ChAT-GFP neurons, the fluorescent intensity of Alexa594 on an individual ChAT-GFP neuron was measured in different parts of the striatum. Results are presented as mean fluorescence of Alexa 594 along the plasma membrane of the cell body (pink line derived from rendering of the cytoplasmic GFP signal in Figure 3B). The results from male and female mice were not different (Supplemental Figure 3A), thus the data were pooled. Average mean fluorescence intensity of NAI-A594 labeled cells in the dorsal striatum was ~20% higher than in ventral striatum (mean fluorescence ±SD = 170.7 ± 57.6 for dorsal, n=52 and 140.9 ± 47.6 for ventral n=47; p<0.05 and n = numbers of ChI neurons, Figure 3C). When the staining of DOR was measured by co-incubating with NAI-A594 plus CTAP, the fluorescent intensity on neurons in the dorsal striatum was reduced to 68% of that observed with NAI-A594 alone (mean fluorescence±SD = 115.5 ± 49.7; n= 41, p<0.0001, Figure 3C). Similarly, the fluorescence of neurons in the ventral striatum was reduced to 61% of NAI-A594 alone (mean fluorescence±SD = 85.8±47.3; n=35, p<0.0001, Figure 3C). Fluorescence of Alexa594-labeled ChAT-GFP neurons in the presence of CTAP was found to be similar to the labeling of the cholinergic neurons from MOR knockout mice (Supplemental Figure 3B) suggesting that the blockade of NAI-A594 labeling at MORs by CTAP was sufficient. Interestingly, when the staining of MOR was determined by NAI-A594 labeling in the presence of SDM25N, the fluorescent intensities of NAI-A594 labeled cells was only 37% and 47% of that observed with NAI-A594 alone in the dorsal and ventral striatum, respectively (mean fluorescence±SD = 63.9±19.5; n=48 for dorsal and 65.7±25.5; n=22 for ventral, p<0.0001 for both Figure 3C). There was a large difference between fluorescent staining of DOR and MOR in dorsal part of the striatum (115.5 ± 49.7, n=41 for DOR *vs.* 63.9±19.5, n=48 for MOR, p<0.0001), while the relative fluorescence of DOR and MOR in the ventral striatum was comparable (85.8±47.3; n=35 for DOR *vs*. 65.7±25.5; n=22 for MOR, Figure 3C). Thus, the results indicate that cholinergic interneurons contain both MORs and DORs and that expression of DORs appears to be higher than MORs in the dorsal striatum but not different from MORs in the ventral striatum. Figure 3D shows the distribution map of NAI-A594 labeled ChAT-GFP positive neurons.

### Inhibition of ChI spontaneous firing activity by MOR and DOR agonists

Cholinergic interneurons are spontaneously active in mouse brain slice preparations with a firing rate of 1-2 Hz (Ponterio et al., 2013). Cell attached extracellular recordings were used to measure the firing activity and the action of MOR and DOR agonists. All recordings were made in the ventral striatum. The average spontaneous firing rate of the GFP-positive neurons was 1.5±0.1 Hz (n=9 cells from 3 male and 3 female mice). Figure 4 shows a representative trace and time-course of firing rate (15-second binning) during agonist and antagonist application. The μ-agonist, DAMGO (1 μM) completely inhibited firing, and this inhibition was reversed following application of CTAP (1 μM).

**Figure 4.**
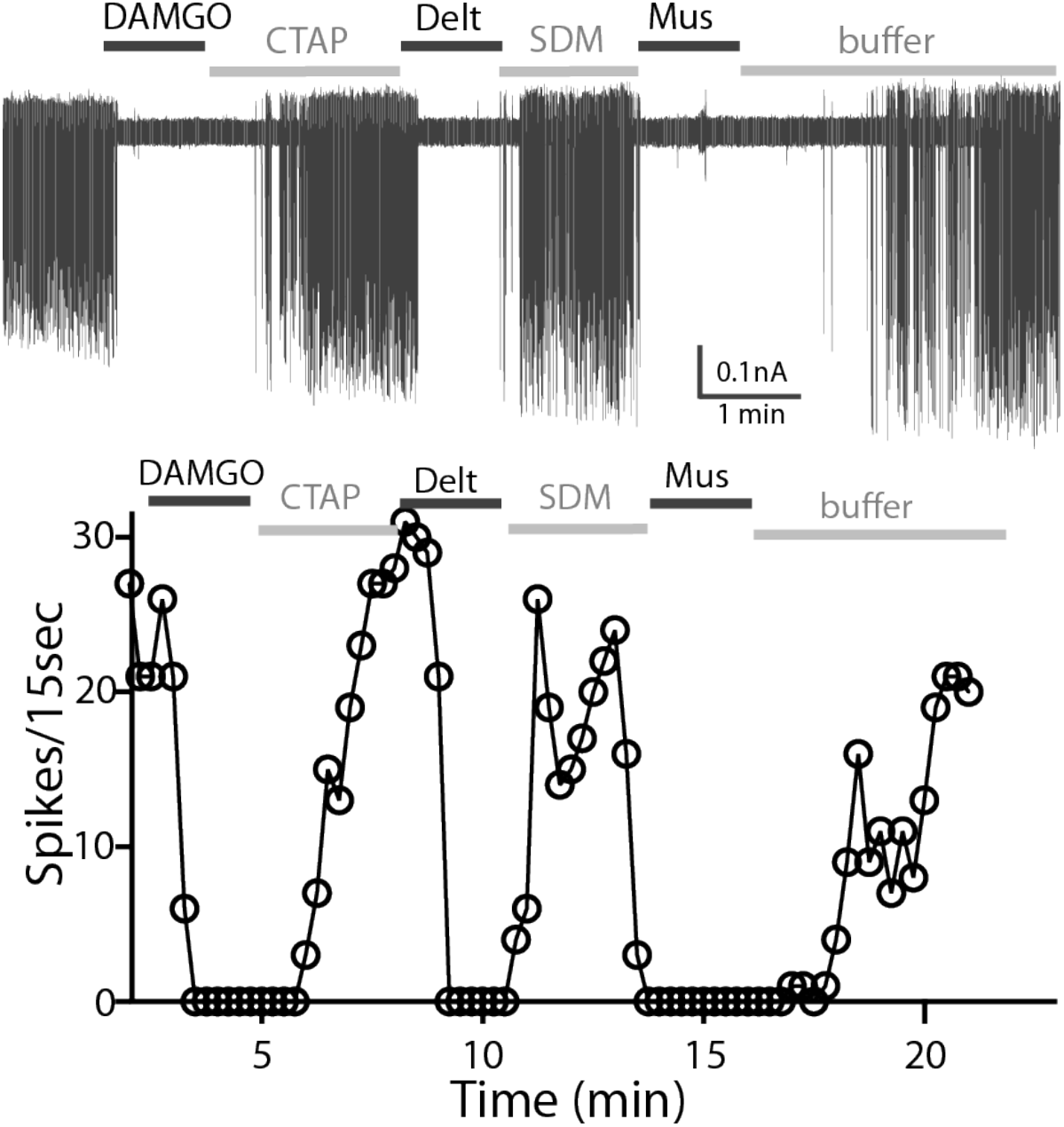
A representative trace of spontaneous firing shows regular spiking at baseline, and a complete stop of firing (100% inhibition) occurs after application of DAMGO (1 μM, 1 min). The firing returns to baseline with application of CTAP (1μM). The subsequent application of deltorphin (1 μM, 1 min) also causes 100% inhibition and this effect is reversed with SDM25N (0.5 μM). Lastly, muscarine (10 μM) causes 100% inhibition of firing. The graph below the trace shows the time course of the experiment plotted against the numbers of spike at 15-second binning.

Subsequent application of the δ-agonist, deltorphin (1 μM) also stopped firing and this could be reversed by SDM25N (0.5 μM, Figure 4A). The inhibition of firing induced by muscarine (10 μM) was included as a control. The results indicate that ChIs co-express functional MORs and DORs.

The effects of opioid agonists on ChI firing inhibition were next studied in neurons from wild type C57/B6 mice. ChIs were identified by the cell morphology and regular firing activity. As in experiments with the slices from the ChAT-GFP mice, neurons in slices from wild type animals were responsive to consecutive application of DAMGO and deltorphin (n=19 cells from 13 mice, data not shown). The capability of each opioid to inhibit ChI spontaneous firing was reported as a percent inhibition calculated by measuring the firing activity before (baseline) and at 1-2 min following agonist application (Table 1).

**Table 1.**
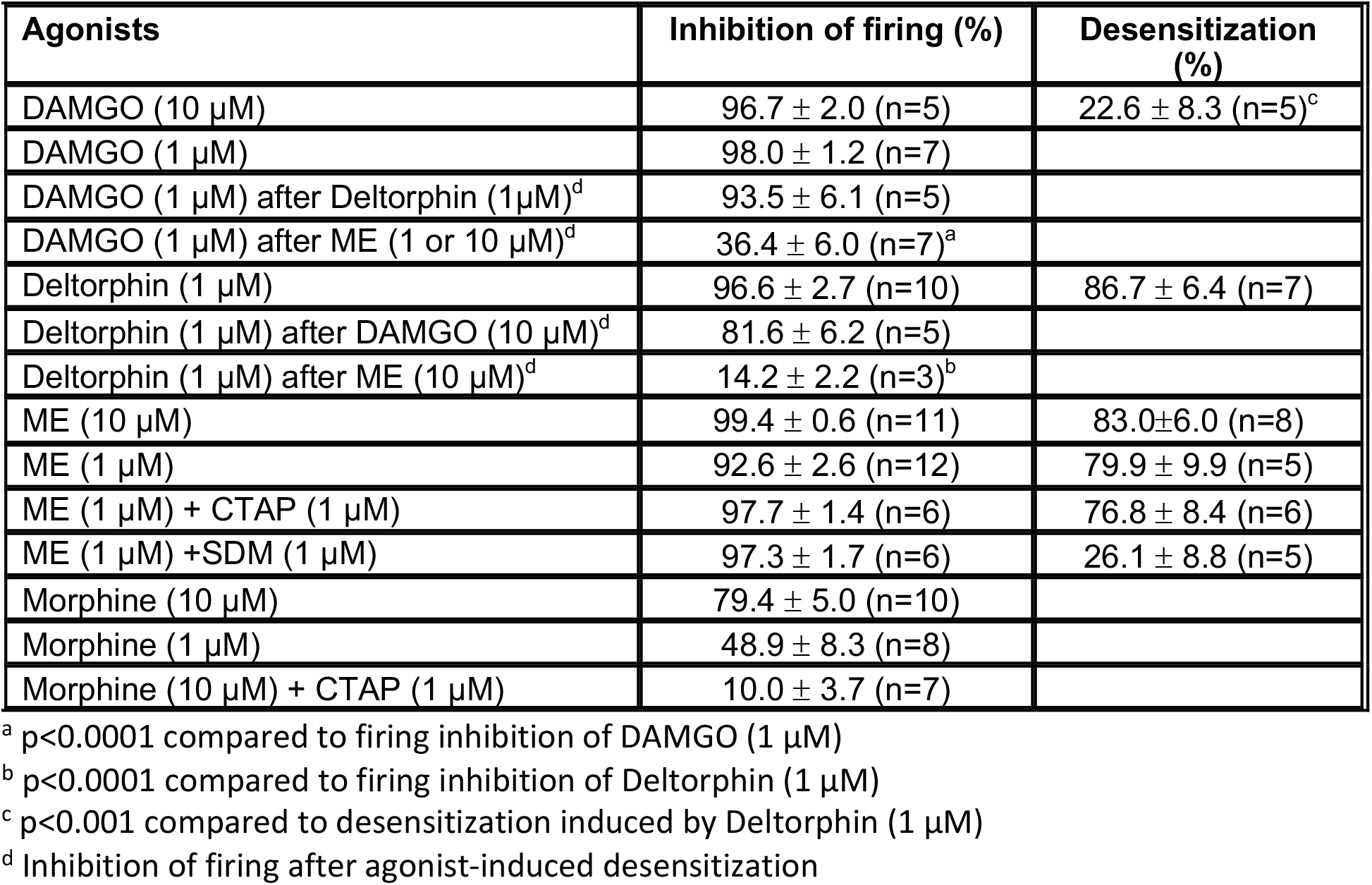
Activation and desensitization by opioids

At baseline, firing frequency of ChIs from wild type mice was similar to that of ChAT-GFP (1.4±0.1 Hz, n=67 cells from 25 mice). Table 1 summarized the inhibition of firing by opioids. DAMGO (1 μM) caused an inhibition of firing activity by 98.0±1.2% (n=7) and increasing the concentration to (10 μM) had no further effect. Deltorphin (1 μM) also caused a total inhibition of firing (96.6±2.7% n=10). Application of a saturating concentration of [Met^5^]enkephalin (ME, 10 μM) resulted in a complete inhibition of the spontaneous firing (99.4±0.6%, n=11) and ME (1 μM) reduced the firing by 92.6±2.6% of baseline (n=12). Because ME acts on MORs and DORs (Gomes et al., 2020), selective antagonists were used to identify the receptor. When CTAP was used to block activity of MORs, ME (1 μM) inhibited the firing activity by 97.7±1.4% (n=6). When DORs were blocked with SDM25N, ME (1 μM) reduced the firing rate by 97.3±1.7% (n=6). Thus, as expected, ME acted on both receptors. Morphine (10 μM) only partially decreased the firing by 79.4±5.0% (n=10) and morphine (1 μM) reduced the firing rate by 48.9 ±8.3% (n=8). In addition, CTAP blocked the inhibition induced by morphine (10.0±3.7% inhibition in 10 μM morphine + 1 μM CTAP, n=7). The results from these experiments provide further evidence of individual activation of MORs and DORs in ChIs by various opioids.

### Receptor desensitization

The application of saturating concentrations of efficacious agonist commonly results in the desensitization of MORs and DORs in many brain areas (Williams et al., 2013; Gendron et al., 2016). The inhibition of firing rate induced by DAMGO and deltorphin was used to determine the extent of MOR and DOR desensitization. Desensitization was determined by calculating ratio of the average firing rate after a 5-min application of agonist compared to the average baseline firing rate (before treatment). Deltorphin (1 μM) initially stopped the firing with a return toward baseline within the 3-5 minutes (Figure 5A, top trace). Thus, deltorphin triggered significant desensitization to 86.7±6.4% (n=7, Table 1). The inhibition of firing induced by DAMGO (10 μM), was however maintained for the entire period of 5-minute application (Figure 5A bottom trace, only 22.6±8.3% of desensitization, n=5, Table 1). The results indicate that DORs are more likely to desensitize than MORs (Figure 5B, p<0.0001).

**Figure 5.**
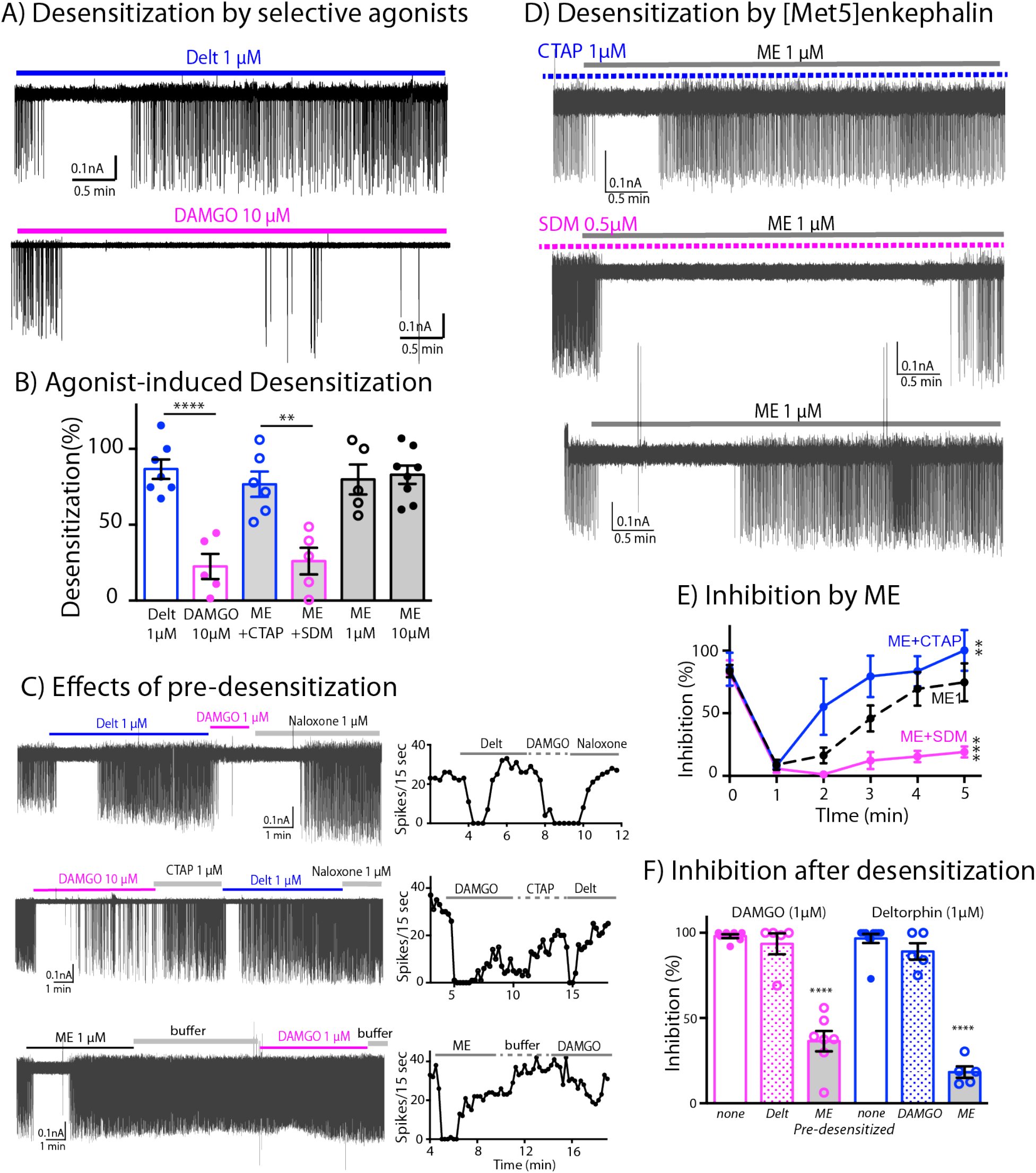
Differential desensitization by selective activation of MOR and DOR. **(A)** *Top* trace shows spontaneous firing of ChI neurons during 5-min application of the DOR selective agonist Deltorphin (1 μM) that caused 100% inhibition of the firing during the first 1-2 minutes and the firing activity gradually returned to nearly baseline during the last 3 minutes of continuous application of the agonist, an indication of desensitization. *Bottom* trace. In contrast, the selective agonist DAMGO caused the persistent inhibition of the firing throughout its 5-min application. **(B)** Summary results of desensitization by Deltorphin, DAMGO, ME (1 and 10 μM) and ME (1 μM) with selective antagonists, CTAP (1 μM) and SDM25N (0.5 μM). **(C)** *Top* trace, desensitization of DORs by deltorphin did not change the ability of DAMGO to stop the firing that was reversed by naloxone. *Middle* trace shows similar experiment in which DAMGO caused little desensitization and subsequent application of deltorphin stopped the firing and rapidly triggered receptor desensitization. *Bottom* trace illustrates the experiment when desensitization by ME decreased the ability of DAMGO to inhibit the firing. Time courses of each experiment plotted as the numbers of spike binned every 15 seconds are shown next to the activity traces. **(D)** Desensitization by ME are very different when co-incubated with CTAP or SDM25N. *Top* trace shows ME desensitization in CTAP developed quickly. *Middle* trace shows no or little desensitization occurred during application of ME plus SDM25N. *Bottom* trace represents desensitization caused by ME alone. **(E)** The graph shows change of % inhibition of firing activity during 5-minute application of ME in different conditions, Two-way ANONA, Sidak’s multiple comparison * p= o.o109 and *** p= 0.0001. **(F)** Summary of firing inhibition shows in *pink*, the ability of DAMGO (1 μM) to inhibit ChI firing activity after pre-desensitized by ME (1 μM) or deltorhine (1 μM) compared to no pre-desensitization, and in *blue*, the firing inhibited by deltorphin (1 μM) after pre-desensitization by ME, DAMGO or none.

To test if this robust desensitization of DORs could affect MOR functions, ChI cells were superfused with deltorphin (1 μM, 5 min) followed by DAMGO (1 μM, Figure 5B top trace). Desensitization caused by deltorphin 1 μM did not change the ability of DAMGO 1 μM to fully inhibit the firing activity of neurons (93.5±6.1% inhibition after deltorphin, n=5 compared to 98.0±1.2% inhibition without deltorphin pre-treatment, n=7, Figure 5F). The reverse experiment where DAMGO (10 μM) was applied first and followed by deltorphin (1 μM) was examined next. In this case, CTAP (1 μM) was used to reverse the firing inhibition of DAMGO before applying deltorphin (1 μM), which was able to inhibit the firing by 89.1±4.8% (n=5, Figure 5C-middle trace and 5F). Again, this inhibition of deltorphin after DAMGO treatment is not statistically different from the inhibition of deltorphin without DAMGO pre-treatment (96.6±2.7%, n=10, un-paired t-test, p=0.1573). The results suggest that MORs and DORs are functionally independent and that there is no major long-lasting cross desensitization between the receptors by selective agonists.

Next, the degree of desensitization at MORs and DORs was examined with the simultaneous activation of both receptors. ME was used in the experiments because it had similarly binding affinity at both receptors (Gomes et al., 2020), and it completely inhibited firing of ChIs (Table 1). Similar to the action of deltorphin, ME (1 μM) plus CTAP fully inhibited the firing that returned to baseline during a 5-minute application (Figure 5D top trace, 76.8±8.4% in Figure 5B, n=6 and Table 1). The desensitization of MORs with ME (1 μM) plus SDM25N resulted in a persistent inhibition (26.1±8.8%, n=5 in Figure 5B and Figure 5D middle trace). Thus, the results indicate that ME will selectively activates and desensitizes MORs or DORs if the other receptor is inactivated by a selective antagonist. In experiments where no antagonists were used, ME (1 μM) caused 79.9±9.9% desensitization (n=5, Figure 5D bottom trace) that was similar to ME (10 μM, 83.0% ±6.0%, n=8). The degree of desensitization caused by ME (1 μM or 10 μM) was not different from that caused by ME plus CTAP or deltorphin (Figure 5B). These results were surprising because the above results predict that activation of MORs by ME should sustain the inhibition of firing activity, since MOR only weakly desensitizes when selectively activated. Time-course analysis of desensitization during 5-minute application of ME in different conditions reveals that desensitization caused by ME activation at both receptors started slower than the desensitization caused by ME plus CTAP (Figure 5E, Two-way ANOVA, p=0.0109 as compared to ME 1 μM).

Conversely, the desensitization by ME alone was faster and more complete than that of ME plus SDM25N (Figure 5E, Two-way ANOVA, p=0.0001, as compared to ME 1 μM). The results suggest that when ME interacts with both MORs and DORs at the same time, MOR desensitization is augmented. To test this assumption, we measured the ability of DAMGO (1μM) to inhibit the firing rate after ME-induced desensitization (Figure 5C bottom trace). The results show a substantial reduction in the inhibition as compared to the control result of DAMGO alone (36.4±6.0% inhibition, n=7 *vs*. 98.0±1.2% of DAMGO alone, n=7, p<0.0001, Figure 5F and Table 1). In addition, the ability of deltorphin (1 μM) to inhibit the firing rate after ME desensitization was also reduced (18.3±3.4%, n=5 inhibition *vs.* 96.6±2.7, n=10, p<0.0001, Figure 5F and Table 1). Thus, ME caused desensitization at both MORs and DORs, and desensitization of MORs could be accelerated by DOR desensitization.

### Receptor internalization

One possible explanation for the increase of MOR desensitization when MOR and DOR are simultaneously activated could be co-internalization induced by ME. To test for this, we used live cell imaging technique to examine agonist-induced internalization of endogenous MORs and DORs that had been labeled with NAI-A594 in brain slice preparations from ChAT-GFP mice. In these experiments, a labeled ChI neuron was first imaged once before agonist application (time = 0 min, Figure 6A-top panel). Then, a saturating concentration of each agonist was applied by superfusion. After 20-minute agonist application followed by an additional 10-min washout, a second image of the same neuron was taken. The control group was treated with only buffer for the same time and images captured at 0 min and 30 min. Fluorescence in the cytoplasm and at the plasma membrane before (F0) and after agonist (F) was measured and used to determine the amount of endocytosis. Figure 6A showed the images of cells before (top panel) and after treatment with different agonists and buffer control (bottom panel). The measurement of fluorescence in the cytoplasm and at the plasma membrane was measured as depicted by an outline derived from GFP signal. (Figure 6B).

**Figure 6.**
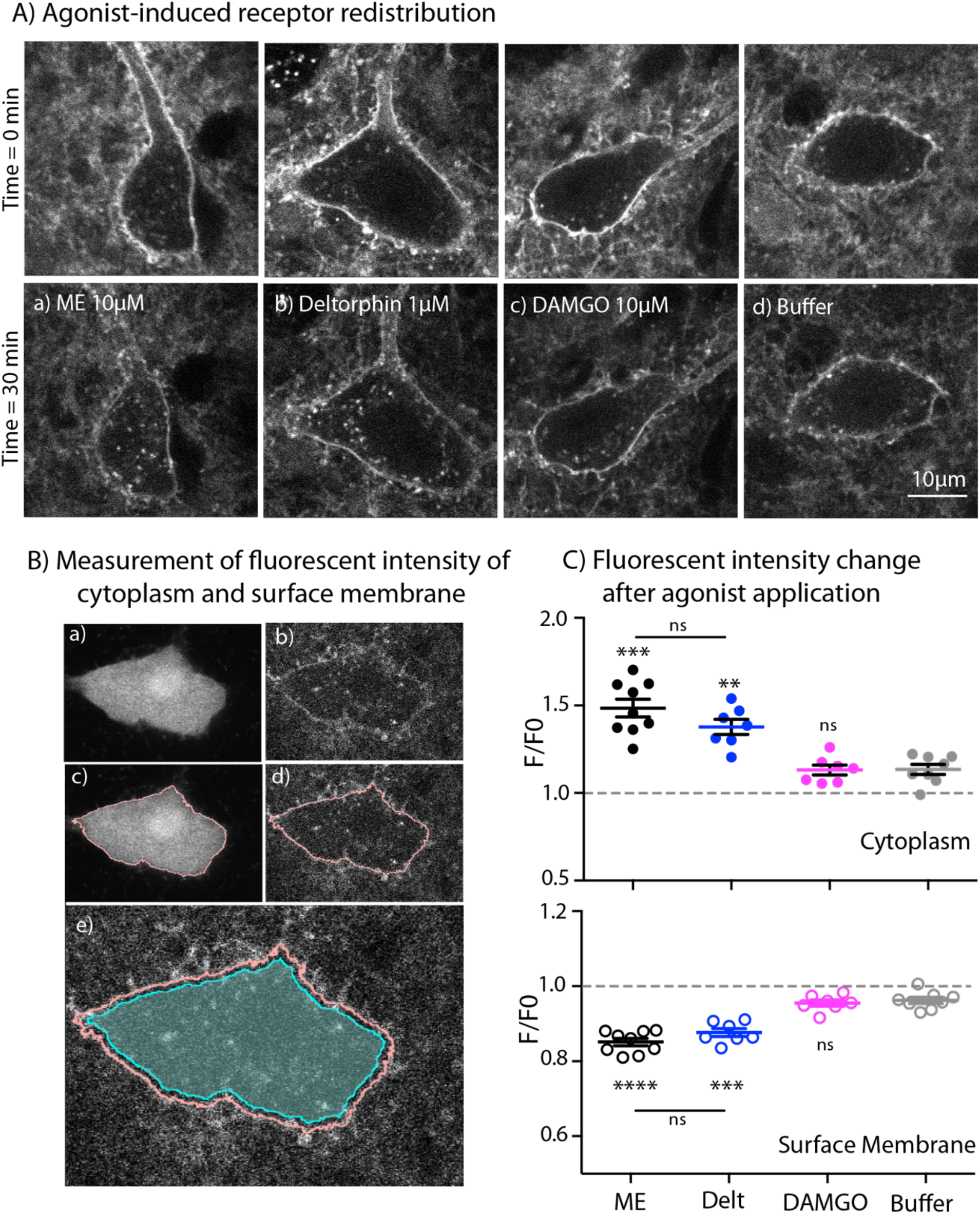
Endocytosis of MOR and DOR. **(A)** Images show receptor re-distribution induced by different agonists. Upper panel shows images taken before agonist application. Each slice was incubated in NAI-A594 100 nM for 1 hour and washed with Kreb’s buffer with continuous bath-perfusion for 10-15 minutes before acquiring the image. Lower panels show images taken after agonist application by bath-perfusion for 20 min and following 10 min of buffer. The control experiment was treated with buffer alone for 30 min. **(B)** Diagram images show measurements of fluorescent intensities along the plasma membrane and inside the cytoplasm using ImageJ. Raw images of one optical slice of a neuron taken simultaneously are shown for GFP (a) and Alexa594 (b). The GFP signal is used to define an outline of plasma membrane (c). This outline is copied to the image of Alexa594 signals (d). The outline of cytoplasm is then dilated 8 pixels (~0.6μm) and the inner line is used to define the area of cytoplasm where internalization taking place (cyan shade area in (e)). Mean fluorescent intensity along the outline of plasma membrane is measured as F-memb (pink line). Mean fluorescent intensity in the area of cytoplasm is measured as F-cyt (cyan shade area). The ratio values of F-memb/(F-memb+F-cyt) and F-cyt/(F-memb+F-cyt) are then calculated for F/F0. F is the ratio value after agonist application and F0 is the value before agonist application. **(C)** Summary of fluorescent intensity changes in cytoplasm (top) and plasma membrane (bottom) of neurons treated with agonists. Data shown as mean±SEM compared to buffer control group, p<0.0001(****), p<0.001(***) and p<0.01(**) compared to control group. ME = [Met^5^] enkephalin, Delt = deltorphin.

ME (10 μM) caused a visible redistribution of the labeled receptors from plasma membrane into the cytoplasm. Fluorescent puncta were found throughout the cell body of the neuron (Figure 6A-a). The F/F0 in the cytoplasm from the ME-treated neurons was significantly higher than F/F0 of cells superfused with buffer alone (1.49±0.05 for ME, n=9 and 1.13±0.03 for control, n=8, p<0.001, Figure 6C, top). Because ME can interact with both MORs and DORs, deltorphin (1 μM) was used to induce endocytosis of DORs. The fluorescent puncta were also clearly observed (Figure 6A-b). The change of fluorescence in the cytoplasm from deltorphin-treated cells was also greater than in the buffer control group (F/F0 = 1.38±0.04 for deltorphin, n=7 *vs*. 1.13±0.03 for control, n=8, p<0.01, Figure 6C, top). Surprisingly, the MOR selective agonist DAMGO (10 μM) barely caused any redistribution of the receptors (Figure 6A, e-h), and the F/F0 of the cytoplasm was the same as that of the buffer control group (1.13±0.03, n=7 DAMGO *vs*. 1.13±0.03 control, n=8, Figure 6C, top).

Next, the fluorescence at the plasma membrane was determined after each treatment. The fluorescence at the plasma membrane is expected to decrease if labeled receptors are internalized. In control experiments the F/F0 on the plasma membrane was 0.96± 0.01 after 30 min (n=8). A value below 1 suggests that a small portion of NAI-A594 may bind reversibly and be removed during washing or some small agonist-independent internalization may occur (Figure 6C, bottom). Following the application of ME or deltorphin, the fluorescence at the plasma membrane significantly decreased (F/F0 = 0.85±0.01 for ME, n=9 and 0.88±0.01 for deltorphin, n=7; p<0.0001 and 0.001, respectively, Figure 6C, bottom). DAMGO caused a slight decrease of the fluorescence ratio at the plasma membrane that was the same as that of the control group (F/F0 =0.96±0.01, n=7, Figure 6C, bottom). The results indicate that endogenous DORs are readily internalized whereas MORs remain mostly on the plasma membrane under these experimental conditions. Importantly, ME-induced receptor internalization was comparable to DOR internalization caused by deltorphin suggesting that ME did not increase MOR endocytosis or cause co-internalization of MOR-DOR.

## Discussion

Co-localization of MOR and DORs has been postulated to exist *in vivo* based on genetically modified receptors in knock-in mice and the use of antibodies (Gendron et al., 2015). In the present work receptor imaging and electrophysiological recordings were used to examine MOR/DOR action and interactions in brain slices from ChAT-GFP and wild type mice. The results demonstrate that while endogenous MORs and DORs are co-expressed on cholinergic interneurons (ChIs), they function independently and do not appear to require stable dimer formation for their activity.

Interactions between two GPCRs *in vivo* could result from physical association of the receptors (aka dimerization) or functional consequence of many possible down-stream events (Lambert and Javitch, 2014). The discovery that MORs and DORs are present and functional in a single ChI neuron in brain slices offers a relevant model system to study these important issues. The results from this study show that a major population of MOR-DOR heterodimers may not stably form, or if physical association does occur, it does not affect the ability of individual receptors to inhibit spontaneous activity of ChIs. In addition, each receptor undergoes distinct regulation processes. DORs desensitize more completely at a faster rate than MORs. Similar results have been reported in studies using cultured neurons from mice with genetic deletions (Walwyn et al., 2009), neuroblastoma SH-SY5Y cells expressing endogenous human MORs and DORs (Prather et al., 1994) and *Xenopus* oocytes exogenously expressing equal amounts of MOR and DOR (Lowe et al., 2002).

When both receptors are concurrently activated with [Met^5^]enkephalin (ME), MOR exhibits an increase in desensitization, which differs from the induction of desensitization at only MORs. The actions of ME on co-expressed MOR-DOR are crucial in understanding functional interactions between these receptors and resultant effects on ChI activity *in vivo*. In conditions where ME independently interacts with each receptor (i.e., in the presence of selective antagonists for MOR and DOR), ME produces similar results to the selective agonists thus underscoring the ability of MORs and DORs to act independently in native environment. The increase in MOR desensitization induced by ME alone where MORs and DORs are simultaneously activated is most likely a heterologous action dependent on downstream processes. It is well known that G protein receptor kinase (GRK) phosphorylation and arrestin-binding are common mechanisms for homologous desensitization of MORs and DORs (Williams et al., 2013; Gendron et al., 2016). Given that ME recruits arrestins to DORs more efficiently than MORs (Gomes et al, 2020), simultaneous co-activation at MOR and DOR by ME may facilitate the interaction between arrestins and MORs. Lowe et al. (2002) reported that in oocyte model, arrestins were the limiting factor for MOR desensitization and increased arrestin expression enhanced the desensitization rate. It will be interesting to know how blockade of GRK phosphorylation or genetic deletion of arrestin might affect the rate and degree of MOR and DOR desensitization in ChIs.

Co-internalization of MORs and DORs would support evidence of physical association of heterodimers. Studies in cultured neurons from mouse dorsal root ganglion (Walwyn et al., 2009) and neurons in the tissue of myenteric plexus from DOR-GFP mice (DiCello et al., 2020) however find no co-internalization induced by a selective agonist at either MORs or DORs. Here, we demonstrate that internalization of MOR and DOR are very different from each other in ChIs. Deltorphin evoked significant DOR redistribution from the plasma membrane into the cytoplasm, but there was no receptor internalization induced by the MOR selective agonist DAMGO. Because receptor internalization by ME was not greater than that induced by deltorphin, the results suggest that the receptors do not traffic together even with simultaneous activation.

One possible criticism of the above analysis would be to propose that NAI-A594 labeled receptors are less likely to be internalized. We think this is not likely to be the case for the following reasons. (1) ME induced the same increases in potassium conductance in A594-labeled and unlabeled locus coeruleus neurons (Arttamangul et al., 2019). (2) Agonist-induced endocytosis of NAI-A594 labeled MORs were internalized in HEK293 cells (Arttamangkul et al., 2019) and in cultured habenular neurons (personal communication with Dr. Damien Jullié, UCSF). (3) With the much smaller molecular size of organic dyes compared to reporter proteins or antibodies used in the study of subcellular localization of receptors (Choquet et al., 2021), it is less likely that A594-tagged receptors would exhibit defective receptor activation. The low signal-to-noise with the use of live brain slices could also limit the detection of low numbers of events and accumulation of the fluorescent signal in the cytoplasm. The trafficking of DORs is mainly targeted to lysosomes (Tsao and von Zastrow, 2000; Scherrer et al., 2006; Pradhan et al., 2009), therefore the fluorescent signal in the cytoplasm would increase with time. It should be noted that although the receptors may be degraded in the lysosomes, the Alexa594 dye will remain fluorescence because of its stability in acidic environment (King et al., 2020). The MORs are known to rapidly recycle back to the membrane surface after endocytosis, especially when the agonist remains bound to the receptors (Yu et al., 2010; Jullié et al., 2019; Kunselman et al., 2019). Because slow wash-out of agonist is common in brain slice preparation, it could be possible that recycling process of MORs in brain slices might occur faster than in cultured cells.

The finding that striatal cholinergic interneurons co-express MORs and DORs presents a new perspective to the role of ChIs in regulating synaptic circuitry in the striatum, particularly after opioid treatment. The present study uncovers distribution of MORs and DORs in ChIs throughout the striatum (see Figure 3D) and the co-localization of MORs and DORs have been functionally studied in the area of ventral striatum. We anticipate that MORs and DORs are also co-expressed on ChIs in the dorsal striatum. Because DORs are more abundant than MORs in the dorsal ChIs (Figure 3C), one would predict that MOR desensitization might be accelerated by co-activation of DORs as found in the ventral ChIs, as the latter would help recruit arrestins, and thus increase the local concentration of arrestins available to MORs. More importantly, functions of co-expressed MOR-DOR may be altered after animals are chronically treated with morphine. A number of studies have reported functional up-regulation of DOR in several brain areas, possibly resulting from an increased receptor expression on the surface membrane (reviewed in Gendron et al., 2014). In DOR-GFP mice, chronic morphine however does not change surface expression of the GFP-tagged DORs on ChIs in the dorsal striatum, and the expression even decreases in the ventral ChIs (Leah et al.,2016).

In summary, the results in this study illustrate co-existence of endogenous MOR and DOR on the surface membrane of ChIs in the striatum. Both receptors function independently and are differently regulated when activated by selective agonists. In spite of their autonomous signaling, the interactions induced by downstream processes such as, phosphorylation and/or arrestin binding, suggest important functional interactions apart from direct receptor association.

## Materials and Methods

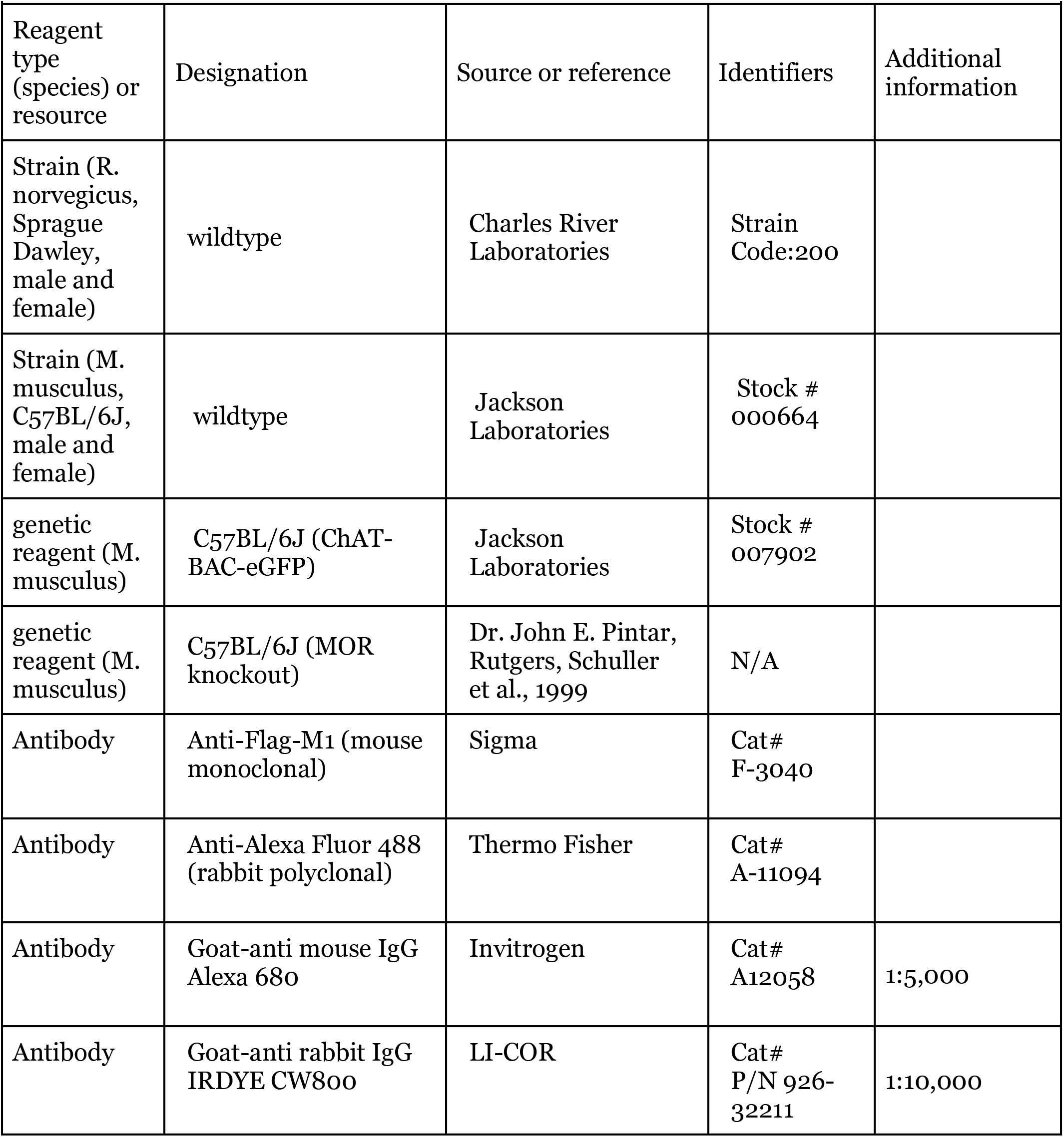
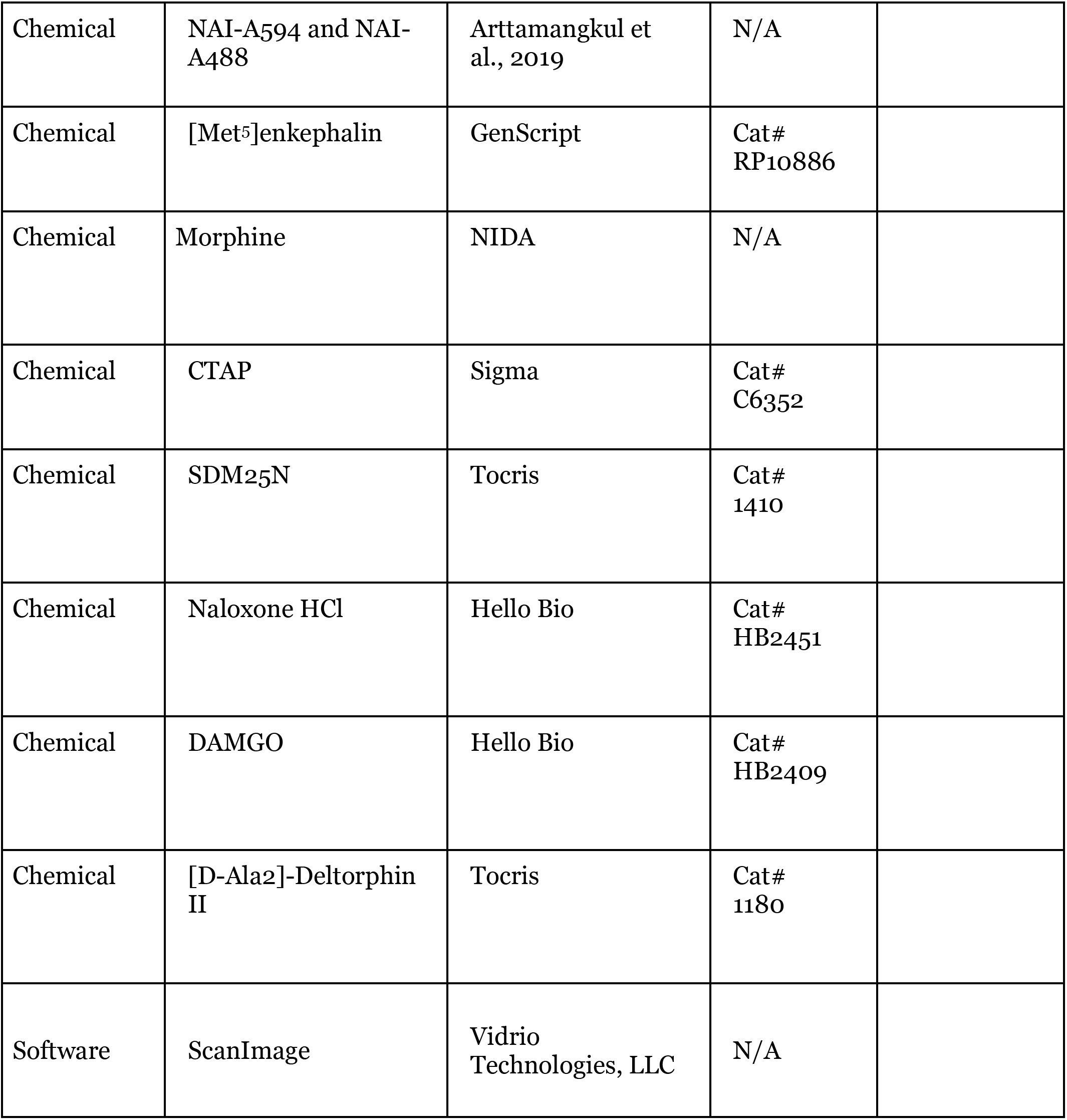

### Drugs and chemicals

NAI-A594 and NAI-A488 were synthesized as previously described (Arttamangkul et al., 2019). 20 nmole of NAI-A594 and NAI-A488 were aliquoted and kept in dried-pellet form at −20°C until used. A stock solution of 100 μM in 2%DMSO-water was made and used within one week. The following reagents were purchased from companies as indicated in parentheses: DAMGO, naloxone and (+)MK-801 (Hello Bio, Princeton, NJ), [DAla^2^]-Deltorphin II and SDM25H HCl (Tocris, Minneapolis, MN), [Met^5^]enkephalin (Genscript, Piscataway, NJ), CTAP (Sigma, St. Louis, MO). Morphine alkaloid was obtained from National Institute on Drug Abuse, Neuroscience Center (Bethesda, MD) and was dissolved in a few drops of 0.1 M HCl and an adjusted volume of water to make a stock solution of 10 mM. All salts to make artificial cerebrospinal fluid (ACSF) used in electrophysiological experiments were purchased from Sigma. Drugs were diluted to the tested concentrations in (ACSF) and applied by bath-perfusion.

### Animals

All animals were handled in accordance with the National Institutes of Health guidelines and with approval from the Institutional Animal Care and Use Committee of OHSU. Mice (21-28 days), both male and female, on a C57BL/6J background were used for all genotypes. ChAT(BAC)-eGFP transgenic mice were purchased from the Jackson Laboratory (Bar Harbor, ME) and the homozygous pairs were bred. Wild type mice were raised from breeders obtained from the Jackson Laboratory. MOR-KO mice were gifted from Dr. John Pintar. Juvenile Sprague-Dawley rats (21-28 days), both male and female, were raised from two breeding pairs that were purchased from Charles River Laboratories (Wilmington, MA).

### Chemical labeling of endogenous MOR and DOR in brain slices

Rats and mice were anaesthetized with isoflurane and brains were removed and placed in warm (30°C) oxygenated ACSF plus MK-801 (0.03 mM). Brain slices were prepared (280 μm thickness) using a vibratome (Leica, Nussloch, Germany). The slices containing striatum between bregma 0.38-1.4 mm of mouse brain and 0.6-2 mm of rat brain were collected and allowed to recover in oxygenated warm ACSF (34°C) containing 10 μM MK-801 for 30 min. Slices were then hemisected. One half was incubated in oxygenated ACSF containing 100 nM NAI-A594 for 1 hr at room temperature. In a separate vial, the other half of brain slice was also incubated for 1 hr at room temperature in the solution of 100 nM NAI-A594 plus CTAP (1 μM) or NAI-A594 plus SDM25N (0.5 μM) to selectively block the labeling of MOR or DOR, respectively.

The labeled slice was then transferred to an upright microscope (Olympus BX51W1, Center Valley, PA) equipped with a custom-built 2-photon apparatus and a 60x water immersion lens (Olympus LUMFI, NA 1.1). Both Alexa594 dye and GFP were excited simultaneously with a high frequency laser beam at 810-nm (Chameleon, Coherent, Inc., Santa Clara, CA). Fluorescence of GFP and Alexa594 were acquired and collected in two separate channels using ScanImage software (Pologruto et al.,2003). Each image acquisition contained 10 optical slices at 0.5 μm thickness. Each optical slice was averaged from 5 consecutive scans of 512×512 pixels. Brain slices were submerged in continuous flow of ACSF at a rate of 1.5-2 ml/min and drugs were applied via superfusion. All experiments were done at 34°C. Macroscopic images of labeled slices were captured with a Macro Zoom Olympus MVX10 microscope and a MV PLAPO2xC, NA 0.5 lens (Olympus). Alexa594 dye was excited with yellow LED (567 nm). The slices were kept under ACSF in a petri dish during data acquisitions. Data were acquired using Q Capture Pro software (Q Imaging, British Columbia, Canada).

### Electrophysiology

Brain slices were prepared as described for the labeling study. A loose cell-attached extracellular recording was done with an Axopatch-1C amplifier in current mode. Recording pipettes (1.7-2.1 MΩ, TW150F-3, World Precision Instruments, Saratosa, FL) were filled with 175 mM NaCl buffered to pH7.4 with 5 mM HEPES. The on-cell pipet resistance varied in the range of 5-15 MΩ. Immediately the cell attaching was formed, spontaneous activity of the neuron was monitored. Only cells that showed steady regular firing at baseline for 3-5 minutes were used in the experiments. Data were collected at sampling rate of 10 KHz, and episode width of 50 second with Axograph X (1.5.4). Drugs were applied by bath perfusion. Bestatin (10 μM, Sigma) and thiorphan (1 μM, Sigma) were included in all peptide solutions to prevent enzymatic degradation.

### Chemical labeling of FDOR and FMOR in HEK293 cells

HEK393 cells stably expressing FlagDOR and FlagMOR were grown and maintained in Dulbecco’s minimal essential media (DMEM, Gibco, Grand Island, NY) containing 10% fetal bovine serum (FBS), Geneticin sulfate (0.5mg/ml, ThermoFisher, Waltham, MA) and antibiotic-antimycotic (1x, ThermoFisher). 24-h before labeling, cells were seeded at 3.0 x 10^5^ cells/well in a 12-well plate. The next day, cells were labeled with 30 or 100 nM NAI-A488 for 30 min at 37°C. Control cells were labeled in the presence of excess 10 μM naloxone to confirm the specificity of labeling. After the labeling period, cells were washed 2 times in PBS + 1μM naloxone, followed by incubation with 10 μM naloxone in PBS containing calcium and magnesium at 37°C for another 10 min. Cells were then placed on ice and washed 2 times in ice-cold PBS plus 1 μM naloxone. The wells were scraped, and cell pellets collected by centrifugation at 4,000 rpm for 4 min at 4°C in an Eppendorf microcentrifuge. Whole cell extracts were produced by resuspending cell pellets in lysis buffer (50 mM Tris pH 7.4, 150 mM NaCl, 1.0% nonidet P-40, 0.1% sodium deoxycholate, 0.5 mM PMSF, 5 μg/ml leupeptin, 1X Halt protease inhibitor cocktail (Thermo Scientific cat. no. 78437)) and nutating for 30 min at 4°C, followed by centrifugation at 14,000 rpm for 20 min at 4°C. The supernatant was collected for further analysis. Protein concentrations were determined using Bio-Rad DC Protein Assay Kit II (cat. no. 5000112) according to the manufacturer’s recommendations. The NAI-A488 labeled cell extracts were then subjected to western blotting.

### Western Blotting

Cell extracts prepared from NAI-A488 labeled cells were analyzed by SDS-PAGE followed by western immunoblotting. 10 μg total protein of each sample was denatured in SDS containing sample buffer (2% SDS, 10% glycerol, 62.5 mM Tris pH 6.8, 0.001% bromophenol blue) in the presence of 0.1 M DTT, at 37°C for 10 min. The samples were then loaded onto 8% SDS-polyacrylamide denaturing gels and run until the dye front just ran off the bottom. The proteins were then electro-transferred to PVDF membranes (Immobilon-FL, cat. no. IPFL00010), blocked in Odyssey Blocking Buffer (LI-COR, cat. no. 927-40000) and probed with two primary antibodies, M1 anti-FLAG MAb (Sigma, cat. no. F3040, used at a 1:600 dilution) and rabbit anti Alexa Fluor 488 (Invitrogen, cat. no. A11094, 1:500), followed by washing and incubation with two secondary antibodies, goat-anti mouse IgG Alexa-680 (Invitrogen, A12058 1:5000 dilution) and goat-anti rabbit IgG IRDye CW 800 (LI-COR cat no. P/N 926-32211, 1:10,000 dilution). Antibodies were diluted in a buffer consisting of 1/10 diluted Odyssey Blocking Buffer, TBS (20 mM Tris pH 7.4, 100 mM NaCl)/1 mM CaCl2 and 0.2% Tween-20. Blots were washed in TBS/1 mM CaCl2/0.3% Tween-20, and final washes were in TBS/1 mM CaCl2. The western blot membranes were scanned using a Sapphire Biomolecular Imager (Azure Biosystems) at 784 nm and 658 nm excitation and images were acquired using Sapphire Capture software (2017).

### On-Cell Western Analysis

96-well optical bottom plates (Thermo Fisher Scientific, cat. no. 165305) were coated with 0.1 mg/ml poly-D-lysine followed by seeding, in triplicate for each condition, 3.0 x 10^4^ DOR or MOR cells per well. 24 hours later, using the same method as above, the cells were labeled with varying concentrations of NAI-A488 (0, 1, 3, 10, 30, 100 nM) and duplicate wells were labeled with each NAI-A488 concentration in the presence of 10 μM naloxone, in order to determine non-specific binding. After the final PBS wash, cells were fixed in 3.7% formaldehyde, 20 min, at room temperature, washed in PBS 3 times and then blocked for 1.5 h, at room temperature, with rocking, in Odyssey Blocking Buffer. The fixed and blocked cells were then incubated overnight at 4°C, on an orbital shaker with primary antibodies (M1, 1:600 and rabbit anti-Alexa Fluor 488, 1:500) diluted in 1:10 Odyssey Blocking Buffer in TBS/1 mM CaCl2. The next morning the wells were washed 5 times, 5 min each in TBS containing 1 mM CaCl2 and 0.05% Tween-20 (OCW wash buffer), followed by incubation in secondary antibodies (Goat-anti mouse IgG-Alexa-680, 1:2500; goat anti-rabbit-IRDye CW 800, 1:5000 diluted in OCW wash buffer) for 1 h, at room temperature while rocking. The wells were then washed 5 times for 5 min in wash buffer, followed by 3 washes for 5 min each in TBS plus 1 mM CaCl2. The wells were air-dried before imaging on a Sapphire Biomolecular Imager using 784 nm and 658 nm excitation. In this experiment, the signal from 784 nm excitation represents opioid receptors modified by NAI-A488, while the 658 nm signal indicates opioid receptor present in each condition. Background values were obtained from cells that were labeled with 100 nM NAI-A488, but that did not receive any primary antibody. Fluorescence intensities were quantitated using AzureSpot Analysis Software (version 2.0.062). All values were background subtracted and the 784 nm signal for each condition was normalized to the 658 nm signal in the same condition. The ratios (784/658) were also calculated for cells labeled with NAI-A488 in the presence of 10 μM naloxone. Prism 6 (GraphPad software, SanDiego, CA) was used to construct concentration dependent curves of non-specific and total labeling and calculate Kd’s.

### Quantitative image analysis of membrane fluorescent intensity

We wrote an ImageJ macro to automate the fluorescent intensity analysis of two-photon microscopy data. The data were collected from two fluorescence emission channels, GFP and Alexa 594. The macro inputs a data stack, first deinterleaving the channels and Gaussian blurring (sigma=1) to smooth the GFP channel’s pixels (Figure 3B). Moments-preserving thresholding is then applied to generate a mask on each slice of the GFP signal. We assigned this GFP mask’s edge corresponding to the shape of neuron (Figure 3B, c). The assignment was verified qualitatively, and the result was well-matched with the signals of NAI-A594 staining on the membrane. Some errors however may occur particularly along dendritic branching planes resulting in truncated masking (down skewing the membrane mean intensity measurement), and that the data in this area would be avoided (see Figure 3C). The mask from each GFP optical slice was then overlaid onto the same optical slice of corresponding Alexa594 channel. The procedure was scanned through every optical section to ensure that the largest area corresponding to the shape of the neuron was included in the analysis. The membrane fluorescent intensity was measured by converting the GFP mask area to a line with a random break introduced along its length. The line traced along the membrane-mask edge and thereby the neuron’s membrane. The mean pixel intensity was collected along the line’s length. Each data stack is analyzed over ten consecutive slices. The mean fluorescence of each optical slice was pooled and computed for the mean membrane fluorescence intensity of a neuron (F-memb). The macro was run without intervention by a researcher who was blinded to the data collection.

### Quantitative image analysis of receptor endocytosis

With the same masking procedure, the analysis comprised two separate operations on the selected mask. The first operation was to determine the mean membrane fluorescent intensity (F-memb) as described earlier. The second operation used an 8-pixel dilation (corresponding to ~0.6 μm) along every point in the membrane mask’s definition to define an area of the neuron’s cytoplasmic portion (Figure 6B). The mean cytoplasmic fluorescent intensity of each optical slice was measured from this area, and all slices’ means were pooled and averaged for the final output of mean cytoplasmic fluorescence (F-cyt) of a neuron.

Receptor endocytosis was determined in two methods. One method calculated the change of cytoplasmic fluorescent intensity from the ratio of F/F0, where F = F-cyt/(F-cyt+F-memb) after drug treatment and F0 = F-cyt/(F-cyt+F-memb) before drug treatment. The other method used F/F0 calculated from membrane fluorescent intensity and F = F-memb/(F-cyt+F-memb) after drug and F0 = F-memb/(F-cyt+F-memb) before drug.

### Statistical analysis

Data are presented as mean±standard error of the mean (SEM) except the data of fluorescent membrane surface intensity, which is shown as mean±standard deviation (SD). The number of sample size is indicated in the figure legends. Statistical analysis and graphs were made with Prism 6 (GraphPad Software, SanDiego, CA). One-way ANOVA with Tukey’s multiple comparison test was used to compare treatment groups and p<0.05 was considered significant. A non-linear saturation of one-site binding curve was constructed from total and non-specific labeling curve and Kd was calculated from curve fitting values using Prism 6.

---

**Supplemental Figure 1.**
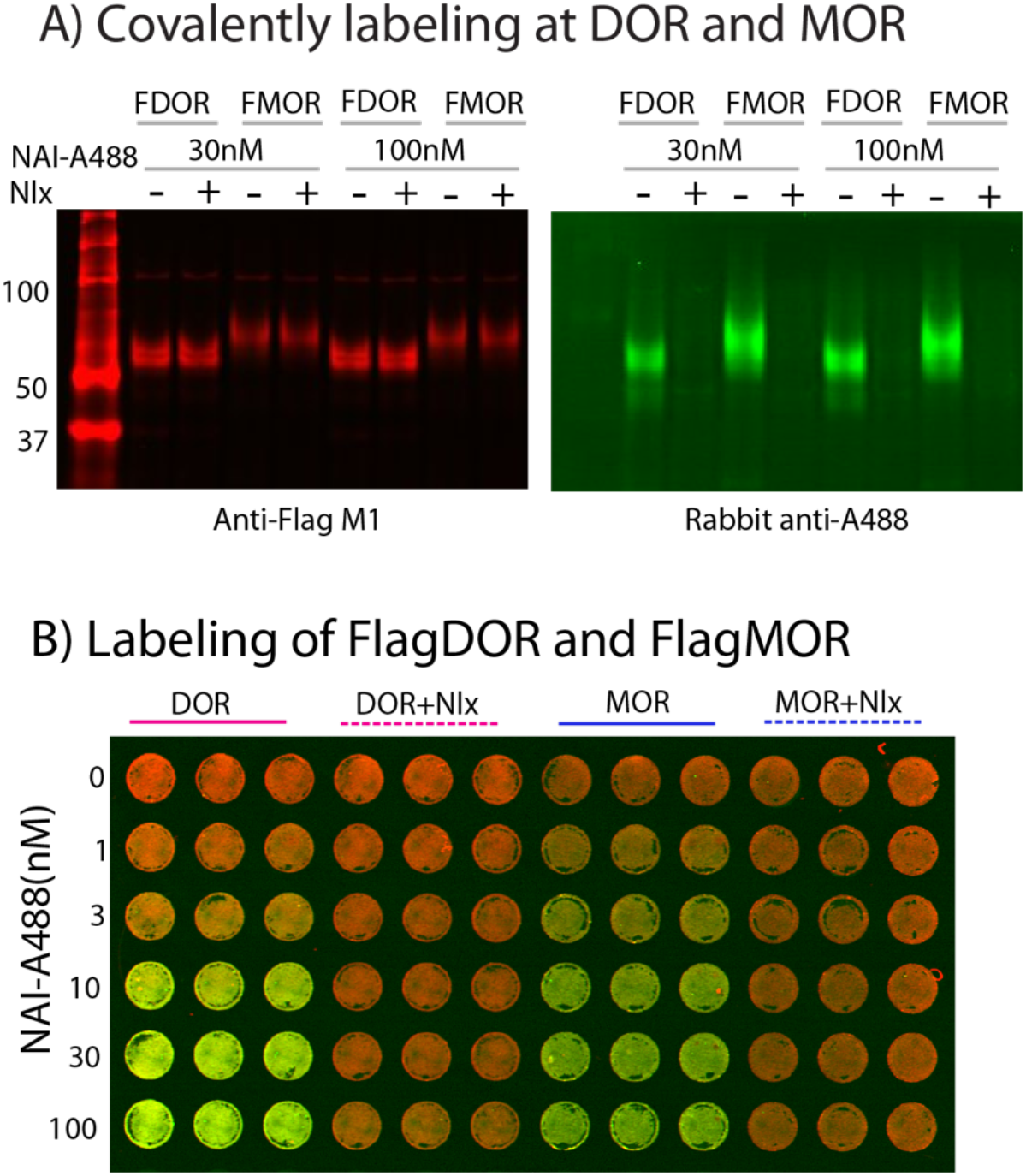
(A) Single channel of the merged image shown in Figure 1 C. (B) On-cell western analysis shows concentration dependence of NAI-A488 labeling FDOR and FMOR cells. The labeling was done in the absence and presence of 10 μM naloxone. Fluorescence intensities were measured with AzureSpot software

**Supplemental Figure 3.**
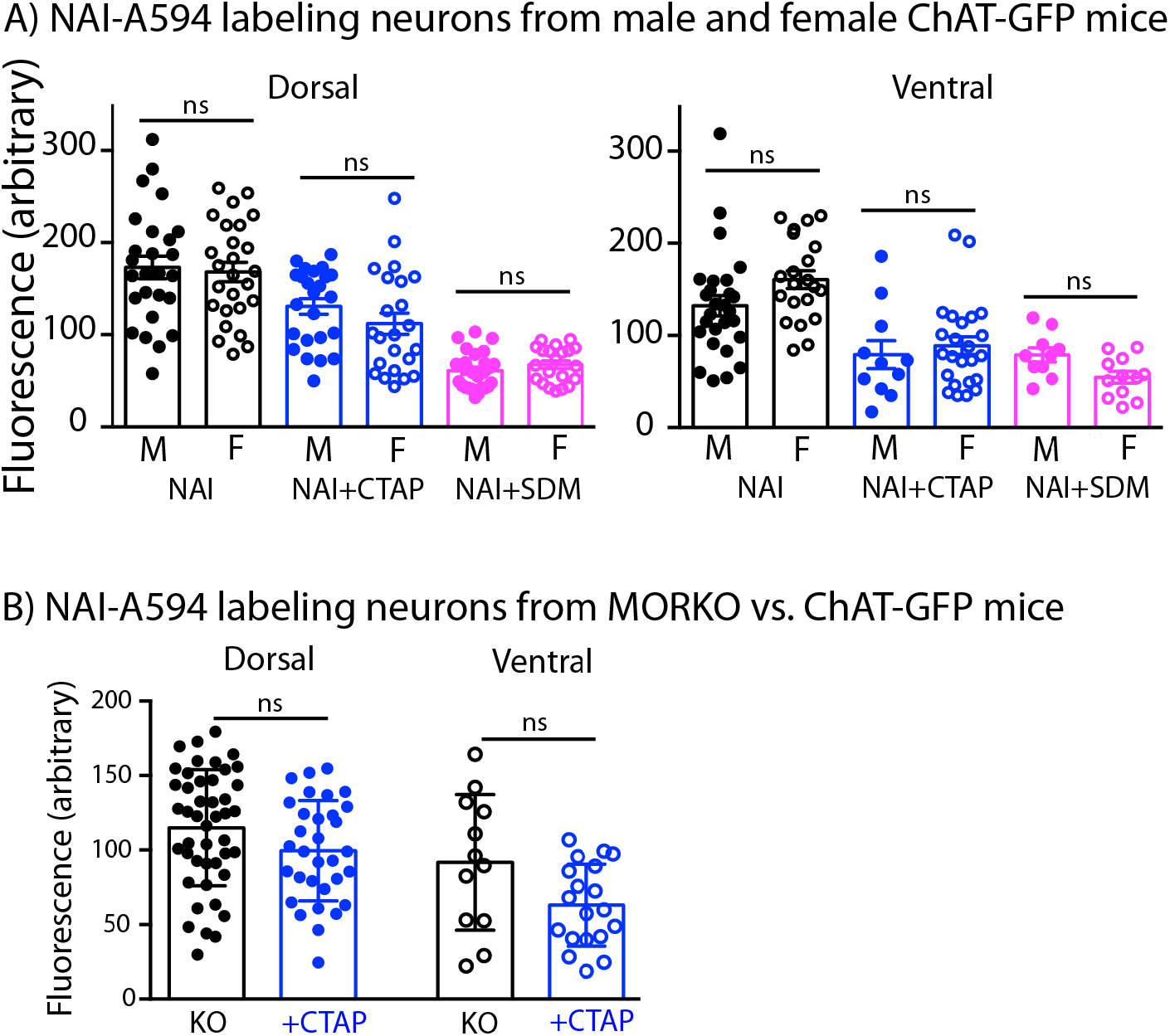
(A)Summary data show that fluorescent intensities in each labeling condition do not differ between male and female mice. Solid dot = male and opened dot = female. Data are shown as mean ± SD. (B)Comparison of fluorescent intensities after labeling of the ChAT-GFP neurons with NAI-A594 plus CTAP and after labeling of ChI neurons from MOR knockout mice with NAI-A594 alone. Because there is no GFP in MOR knockout cells, both measurements were done manually by drawing the outline corresponding to the Alexa594 signal. Data are shown as mean ± SD.

## Acknowledgments

The authors thank Dr. John T. Williams for his constructive advice on the project and manuscript and Dr. Marina E. Wolf for her critical comment on the manuscript. We also thank Drs. Aaron Nilsen and Victoria S. Halls at the Medicinal Chemistry Core, OHSU for the synthesis service of fluorescent NAI compounds and Dr. Kenner Rice at Drug Design and Synthesis Section, NIDA and NIAAA for providing of naltrexamine. The work is funded by grants DA048136 from the National Institutes of Health.

## Author Contributions

S.A. initiated the project, performed experiments, analyzed data and wrote the manuscript. EJP and DLF designed the biochemical experiments of western blot and on cell western analysis. EJP performed experiments and analyzed data. JC wrote a macro program for batch analysis of fluorescent measurement and performed data analysis. All authors read and edited the manuscript.

